# Herpes Simplex Virus Co-Infection Facilitates Rolling Circle Replication of the Adeno-Associated Virus Genome

**DOI:** 10.1101/2020.12.23.424160

**Authors:** Anita F. Meier, Kurt Tobler, Remo Leisi, Anouk Lkharrazi, Carlos Ros, Cornel Fraefel

**Author notes:** These authors contributed equally to this work (Anita F. Meier, Kurt Tobler). Cornel Fraefel, Winterthurerstrasse 266a, 8057 Zurich, Switzerland.

## Abstract

Adeno-associated virus (AAV) genome replication only occurs in the presence of a co-infecting helper virus such as adenovirus type 5 (AdV5) or herpes simplex virus type 1 (HSV-1). AdV5-supported replication of the AAV genome has been described to occur in a strand-displacement rolling hairpin replication (RHR) mechanism initiated at the AAV 3’ inverted terminal repeat (ITR) end. It has been assumed that the same mechanism applies to HSV-1-supported AAV genome replication. Using nanopore sequencing as a novel, high-throughput approach to study viral genome replication we demonstrate the formation of double-stranded head-to-tail concatemers of AAV genomes in the presence of HSV-1, thus providing evidence for an unequivocal rolling circle replication (RCR) mechanism. This stands in contrast to the textbook model of AAV genome replication when HSV-1 is the helper virus.

**SIGNIFICANCE:** Efficient adeno-associated virus (AAV) replication requires the presence of helper factors, which can be provided by co-infecting helper viruses such as adenoviruses or herpesviruses. AAV replication has been described to occur as a rolling hairpin replication mechanism. However, we show that during a herpes simplex virus type 1 (HSV-1) supported replication, AAV rolling circle-like replication intermediates are formed. Thus, this study stands in contrast to the textbook model of AAV genome replication. Additionally, we introduce nanopore sequencing as a novel, high-throughput approach to study viral genome replication in unprecedented detail.

## INTRODUCTION

Adeno-associated virus (AAV) is a small, non-pathogenic *Dependoparvovirus* and is predominantly known for its application in gene therapy (1, 2). The wildtype AAV genome consists of a single-stranded DNA (ss DNA) with a length of 4.7 kb containing a *rep* and a *cap* coding region flanked by 145 nucleotides long inverted terminal repeat (ITR) sequences (3, 4). AAV can only replicate in presence of helper factors, which can be provided by co-infecting helper viruses such as adenoviruses (AdV) or herpesviruses (5, 6). Helper factors are required to transactivate transcription of AAV genes and support AAV genome replication. It was suggested that concatemeric AAV DNA isolated from AAV2/ AdV5 co-infected cells is organized as alternating plus and minus strands (7). However, that study did not provide definite proof of such concatemers. Nevertheless, that observation together with preceding studies on the replication of autonomous parvoviruses and the proposal of a novel replication model for linear double-stranded DNA (ds DNA) (8) led to the currently widely accepted model of the rolling hairpin replication (RHR) of the AAV genome (8–13). According to that model the RHR of the AAV genome is initiated at the palindromic terminal ITR sequences, which are capable of self-annealing into double hairpin structures (3) and thus providing the 3’-OH primer. Extension of the annealed 3’ end leads to the generation of a duplex structure, which is covalently closed at one end. To resolve the closed end, a nick is introduced into one strand of the covalently closed ITR sequence at the terminal resolution site (*trs*) by AAV Rep68/78 (14). The newly generated 3’-OH primer is then used to fill in the remaining gap to form a full-length linear open duplex configuration. Following denaturation of the ends and the subsequent re-annealing into double hairpin structures, a new 3’-OH primer is formed, and, upon strand-displacement, the progeny genome is released.

Several lines of evidence suggest that the described model is valid for AdV-supported AAV genome replication: (i) The predicted covalently closed double-stranded (ds) monomer structures have been isolated from AAV-infected cells (7, 12). (ii) It was shown that the palindromic ITR-sequences behave like origins of DNA replication (12, 15, 16). (iii) Two different ITR orientations (flip and flop) have been observed and their relative orientation to each other was found to be independent (3, 17). The observation of two ITR orientations is in accordance with the model as it predicts that the ITRs are inverted relative to each other during each round of replication. (iv) Site-specific nicking of the *trs* was observed (14, 18, 19). (v) It has been suggested that concatemers isolated from AAV/ AdV co-infected cells consisted of alternating plus and minus strand AAV genomes (7). Such concatemers are believed to arise in the absence of *trs*-nicking. It has been generally accepted that herpes simplex virus type 1 (HSV-1)-supported AAV genome replication occurs in the same manner as described for AdV-supported replication.

The widely accepted model for HSV-1 genome replication is that HSV-1 DNA circularizes upon entry, followed by an origin-dependent bidirectional replication, which then switches to a rolling circle replication (RCR) mechanism (20, 21). It has further been shown that addition of HSV-1 proteins essential for genome replication to non-viral plasmid DNA mediates amplification of the plasmid and leads to the formation of head-to-tail concatemers (22, 23). Since the AAV genome was found to circularize within the host cell (24, 25), which is a prerequisite for RCR, we assessed if HSV-1 mediated AAV genome replication occurs in an RCR mechanism. We explored AAV genome replication intermediates using nanopore sequencing and Southern analysis. The nanopore sequencing technology provides the unique feature of extremely long reads and therefore enables the analysis of single replication intermediate molecules. As hypothesized, we found head-to-tail concatemers, which cannot be explained by the current model of AAV genome replication. Therefore, we suggest RCR as a novel replication model for the AAV genome amplification in presence of HSV-1 helper factors. In addition, we present nanopore sequencing as a novel, high-throughput approach to investigate the mechanism of viral genome replication. Nanopore sequencing has previously been applied to analyze the integrity of genomes of recombinant AAV vector stocks (26). However, we are unaware of any studies using nanopore sequencing for the unbiased identification of genome replication products and the assessment of viral genome replication mechanisms in infected cells.

## RESULTS

### Categories of replication intermediates observed with nanopore sequencing

AAV genome replication may result in intermediate products that are not all detected in bulk approaches such as Southern blotting. Nanopore sequencing provides unique features, which are useful for the assessment of diverse nucleic acid populations. Sequencing length is (theoretically) unlimited and no fragmentation of sample material is required. We employed nanopore sequencing in order to identify and quantify the full range of AAV2 DNA replication intermediates in a high-throughput manner. For this, we extracted extrachromosomal DNA from BJ cells infected with non-replicating (single-infection) or replicating AAV2 (HSV-1 co-infection) using the Hirt protocol and sequenced the isolated DNA with nanopore technology (**Figure 1a**). Notably, extracted DNA was not linearized prior to sequencing. Since adapter ligation for nanopore sequencing requires linear genomes, circular genomes are automatically excluded from our analysis.

**Figure 1.**
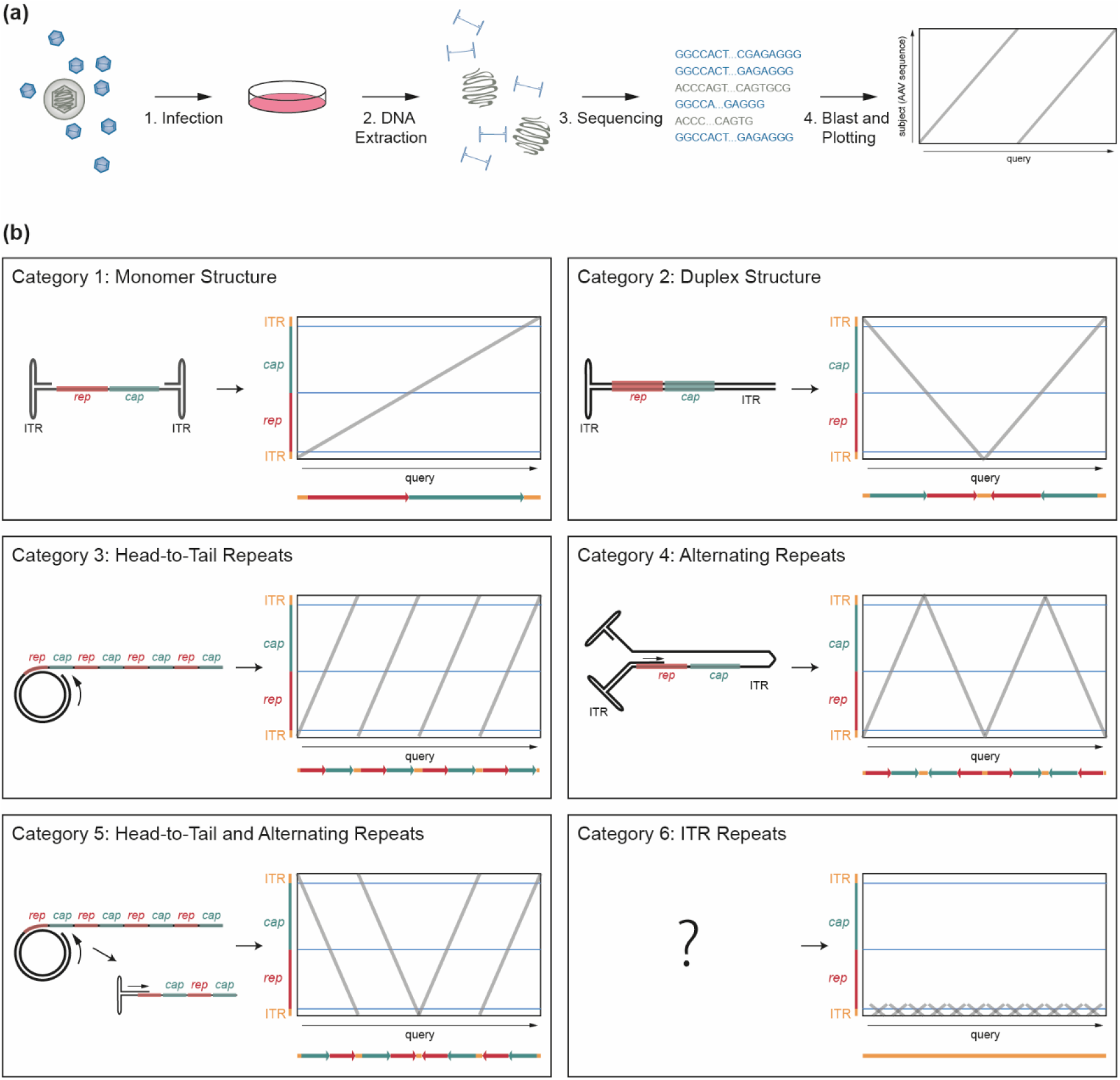
Experimental setup and definition of categories of AAV2 DNA replication products. (a) Schematic representation of the experimental setup. (b) Schematic representation of the defined categories with the predicted AAV DNA structure on the left and the corresponding theoretical dot plot on the right. Dot plots were used to visualize the AAV sequence from isolated genomes with the read (query) on the x-axis and the location on the AAV genome on the y-axis. For further clarification, the read sequence is shown below the dot plot. The AAV ITR-ends (orange) and the genome coding regions *rep* (red) and *cap* (green) are indicated.

To facilitate the comparison of the heterogenous DNA replication intermediates, AAV2 DNA reads were depicted as a function of the included genomic domains. Based on the characteristic patterns obtained, six categories of sequencing reads were defined as shown in **Figure 1b**. Category 1 reads appear as a single straight line crossing over to the other side of the graph from bottom to top (positive polarity) or top to bottom (negative polarity). Such reads represent single AAV2 genomes. Category 2 reads appear in a “V” or an “A” shape on the dot plot. These reads correspond to the duplex form of the AAV2 genome, two ssDNA genomes which are inverted relative to each other and covalently linked on one side as shown below the dot plot. Such structures would arise from single-to-double-strand conversion. Category 3 reads appear as parallel lines crossing the plot that represent multiple direct genome repetitions (head-to-tail concatemers) and would be expected from a rolling circle replication (RCR) mechanism. Category 4 reads appear as zigzag lines in the dot plot that represent multiple copies of the AAV2 genome with alternating orientation and would be expected from a rolling hairpin replication (RHR) mechanism, where *trs*-nicking did not occur. Category 5 reads appear as parallel lines broken up by “V” or “A” shaped structures that represent sequences combining both head-to-tail as well as alternating repeats. We suggest that those concatemers are generated by RCR followed by the re-annealing of the 3’-OH ITR and subsequent second-strand synthesis of the head-to-tail concatemers, resulting in the inversion of the RCR structure. Category 6 reads represent multiple ITR repetitions (no *rep-* and *cap-* sequences). Sequences that could not be assigned to any of the six categories were grouped in category 7. Many of those reads appear as defective, incomplete or shortened genomes (see **Supplementary Figure S1** for representative dot plots of reads assigned to category 7). It is unclear, how such reads were generated but their pattern suggests a defective replication mechanism.

### Nanopore sequence analysis of replicating AAV2 reveals RCR-like intermediates at high frequency

We sequenced extrachromosomal DNA isolated from AAV2 and HSV-1 co-infected cells and found a wide range of DNA replication intermediates. These reads were categorized as described in the section above (**Figure 1b**). We categorized 200 randomly selected reads per sample or all reads of the sample if the number was below 200. In **Figure 2a-f** we show two representative dot plots of individual reads of each category; the frequency of these reads is shown in **Table 1**. According to Oxford Nanopore Technology, our adapter-ligation approach should exclude any ss DNA for sequencing. Since encapsidated AAV genomes are present as ss DNA of positive or negative orientation in equal amounts (27), we assume that strand-annealing occurred prior to library preparation and thus we were able to detect monomer AAV2 genomes.

**Table 1.** Read analysis data from genomes isolated from purified AAV2 virus stock or AAV2 single- or HSV-1 co-infected BJ cells at 12 hpi.

| Category: |  | 1 | 2 | 3 | 4 | 5 | 6 | 7 |  |
| --- | --- | --- | --- | --- | --- | --- | --- | --- | --- |
| Sample |  | Monomer | Duplex | Head-to-Tail Repeats | Alternating Repeats | Head-to-Tail and Alternating Repeats | ITR Repeats | Others |  |
|  | AAV gcp/ cell | ratio | ratio | ratio | ratio | ratio | ratio | ratio | total reads |
| AAV2 stock | NA | 0.830 | 0.040 | 0.005 | 0.005 | 0.000 | 0.065 | 0.055 | 200 |
| AAV2 stock | NA | 0.810 | 0.020 | 0.030 | 0.000 | 0.000 | 0.065 | 0.075 | 200 |
| AAV2 | 20k | 0.620 | 0.080 | 0.015 | 0.005 | 0.000 | 0.190 | 0.090 | 200 |
| AAV2/ HSV-1 | 20k | 0.340 | 0.170 | 0.170 | 0.035 | 0.060 | 0.100 | 0.125 | 200 |
| AAV2 | 500 | 0.250 | 0.333 | 0.000 | 0.000 | 0.000 | 0.167 | 0.250 | 12 |
| AAV2/ HSV-1 | 500 | 0.175 | 0.385 | 0.215 | 0.010 | 0.075 | 0.010 | 0.130 | 200 |

**Figure 2.**
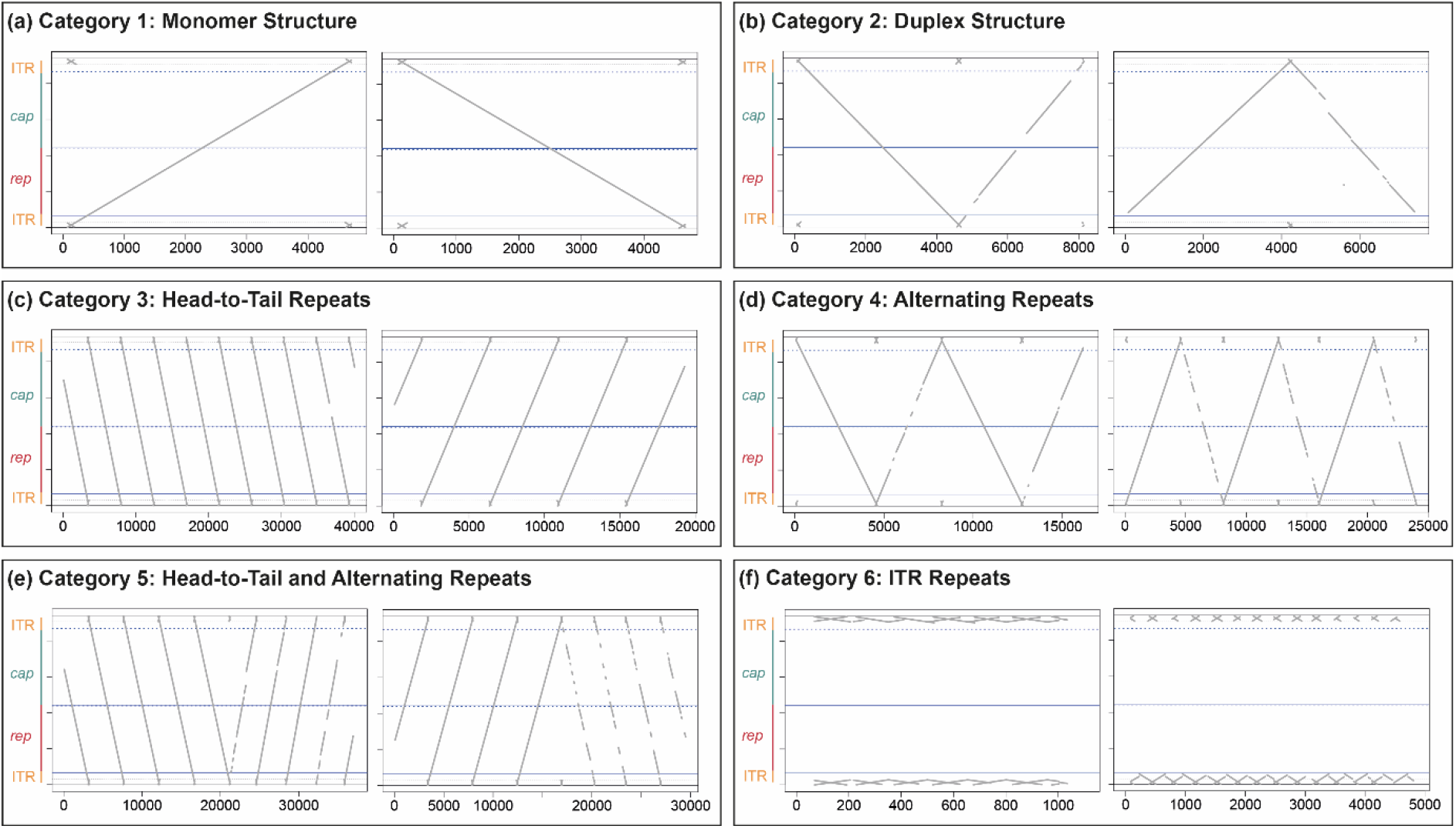
AAV2 replication intermediates. (a)-(f) Dot plots of two representative individual reads per category from AAV2/ HSV-1 co-infected BJ cells are shown. Extrachromosomal DNA was isolated at 12 hpi from BJ cells infected with AAV2 (gcp/ cell = 20’000) and HSV-1 (pfu/ cell = 1).

In samples of extrachromosomal DNA isolated from AAV2 infected BJ cells, the AAV2 DNA was mainly present as monomeric AAV2 genomes (category 1) (62% at AAV2 genome containing particles (gcp)/ cell = 20’000) (**Table 1**). 8% of AAV2 reads were present in the duplex form (category 2). Most of the remaining reads were assigned to category 6 (19%), which contains reads with ITR-repeats, or to category 7 (9%), which contains all other structures. These results were in line with our expectation of predominant monomeric AAV2 genomes and the occasional conversion of ss to the ds DNA duplex structure.

In contrast, AAV2 reads from HSV-1 co-infected cells showed a dramatic change in the observed genome structures (**Figure 2, Table 1**). The ratio of monomeric AAV2 genome reads (category 1) was reduced to 34% for AAV2 gcp/ cell = 20’000 or 17.5% for AAV2 gcp/ cell = 500 (**Table 1, Figure 2a**). Formation of duplex structures (category 2) went up to 17% or 38.5% for AAV2 gcp/ cell = 20’000 or for AAV2 gcp/ cell = 500, respectively (**Table 1, Figure 2b**). The most remarkable result is that HSV-1 co-infection led to the appearance of head-to-tail concatemers (category 3) at 17% or 21.5% (for AAV2 gcp/ cell = 20’000 or for AAV2 gcp/ cell = 500, respectively) (**Table 1, Figure 2c**). We also observed concatemers of alternating orientation (category 4; head-to-head) that would be expected to result from a rolling hairpin replication mechanism upon omission of *trs*-nicking (3.5% or 1% for AAV2 gcp/ cell = 20’000 or for AAV2 gcp/ cell = 500, respectively) (**Table 1, Figure 2d**). However, these reads were far less abundant than the head-to-tail concatemers. Interestingly, structures presumably formed upon second-strand synthesis of an RCR product (category 5) were more abundant than concatemers with alternating genome orientation (category 4). 6% and 7.5% of reads from samples with AAV2 gcp/ cell = 20’000 or for AAV2 gcp/ cell = 500, respectively, were assigned to category 5 (**Table 1, Figure 2e**). Interestingly, in the presence and absence of HSV-1, we identified a high abundance of reads which contain multiple repetitions of ITR sequences classified as category 6 (**Figure 2f**). Those structures were present in up to 19% of all reads and were generally more abundant in absence of HSV-1 (**Table 1**). Furthermore, we also identified those ITR-repeat containing reads in the virus stock samples with an abundance of 6.5% (**Table 1**). The source of those structures remains to be elucidated. AAV DNA, isolated from BJ cells infected with the replication-competent AAV2 variant AAV201, shows a comparable abundance and distribution of structures as seen in BJ cells infected with wtAAV2 (**Supplementary Table S1**).

Quantification of the AAV2 stock reads showed that the vast majority (83 and 81%) were present as monomeric DNA of approximately 4.7 kb with either positive or negative polarity (category 1) (**Table 1**). The remaining population of reads were assigned to category 2 (duplex) (4 and 2%), category 3 (head-to-tail repeats) (0.5 and 3%), category 6 (ITR-repeats) (6.5%) or category 7 (other structures) (5.5 and 7.5%) (**Table 1**). The maximum size of most of those rather unexpected reads was around 5 kb, corresponding with the maximum packaging size of AAV (28). This indicates that a considerable proportion of about 10-15% of viral particles contained defective genomes. Overall, these sequencing results were in line with our predictions, since we reasoned that most of the viral particles from the AAV2 stock sample contain a monomeric AAV2 genome.

In conclusion, monomeric AAV2 genomes were highly abundant in all samples. Complete duplex structures were observed only in DNA samples isolated from infected cells but not from AAV2 stock samples. Furthermore, head-to-tail concatemers were identified at high rates in samples from cells co-infected with AAV2 and HSV-1. Concatemers with a head-to-head structure expected from an RHR mechanism lacking *trs*-nicking were found only at very low rates. Therefore, we conclude that HSV-1 co-infection induces the formation of RCR-like head-to-tail concatemers of the AAV2 genome.

### Southern analysis of replicating AAV2 DNA reveals RCR products

We further assessed the structure of AAV2 genomes isolated from BJ cells infected with AAV2 or co-infected with AAV2 and HSV-1 by Southern analysis. Hirt-DNA extraction of cells infected with AAV2 only resulted in the appearance of a single band with a migration corresponding to about 3 kb of ds DNA (**Figure 3b**). This band represents the monomeric ss AAV2 DNA, as it disappeared upon mung bean nuclease (MBN) treatment, which specifically degrades ss DNA-containing fragments **(Figure 3b)**. The same band was found in samples isolated from AAV2 and HSV-1 co-infected cells, which also disappeared upon MBN treatment (**Figure 3b**). HSV-1 co-infection led to the appearance of a MBN-resistant 4.7 kb band, representing ds AAV2 DNA (duplex) structures. In addition, higher molecular weight bands were observed, which represent concatemers (**Figure 3b**). Of note, higher molecular weight concatemers are underrepresented on the blot because these molecules transfer much less efficiently from the agarose gel onto the nylon membrane than smaller molecules. To reveal the nature of those concatemers, the samples were treated with *Hin*dIII, which cuts ds AAV2 genomes at nucleotide position 1882, within the *rep* sequence (**Figure 3a**). *Hin*dIII digestion of monomeric ds AAV2 DNA is expected to yield fragments of approximately 1.9 kb and 2.8 kb (**Figure 3a**). Since the Southern probe hybridizes to a *rep*-sequence within the 1.9 kb fragment, only this band can be observed. *Hin*dIII digestion of head-to-tail concatemers would result in genome-sized 4.7 kb fragments (**Figure 3a**). Digestion of head-to-head concatemers would result in a 3.8 kb and a 5.6 kb fragment but the hybridization probe only binds to the 3.8 kb fragment. *Hin*dIII does not digest ss DNA. *Hin*dIII treatment resulted in the disappearance of the concatemers observed in the samples from cells co-infected with AAV2 and HSV-1. Instead, a strong band of approximately 4.7 kb as well as two bands of approximately 3.8 kb and 1.9 kb appeared (**Figure 3b**). Since the 4.7 kb band was much stronger than the 3.8 kb band, this demonstrates that AAV2 genome replication intermediates isolated from cells co-infected with HSV-1 are present predominantly as head-to-tail concatemers of ds DNA. These results are in line with our sequencing results described above. Comparable results were obtained when Southern analysis was performed using extrachromosomal DNA extracted from AAV2 and HSV-1 co-infected 293T and Hela cells (**Supplementary Figure S2**).

**Figure 3.**
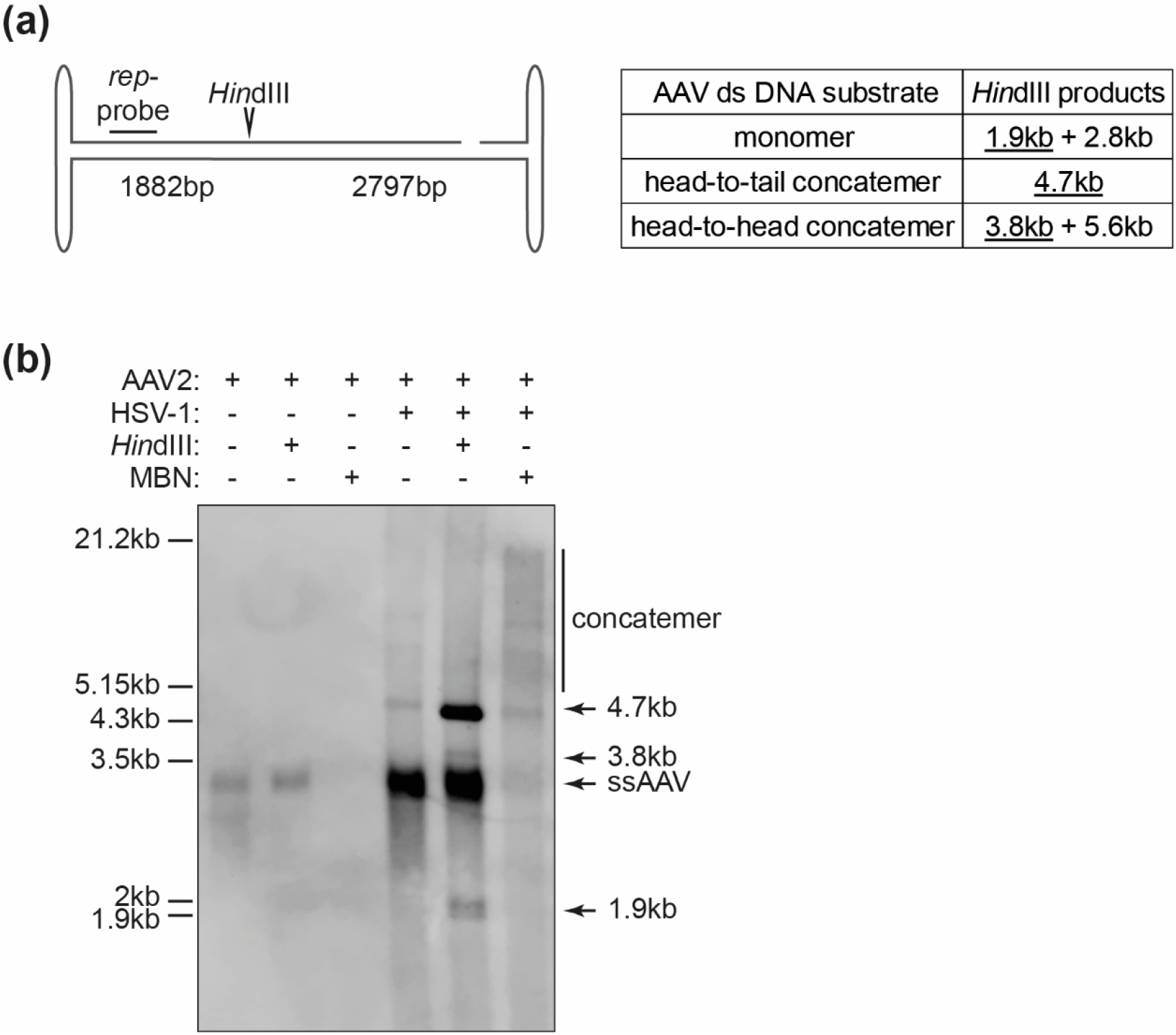
Southern analysis of AAV2 DNA. (a) Schematic representation of the AAV2 genome with the *Hin*dIII restriction site and the *rep-*probe binding site indicated. The table on the right shows the expected sizes of the fragments upon *Hin*dIII treatment with the fragment containing the *rep-*probe binding site <u>underlined</u>. (b) Southern blot of Hirt DNA extracted at 12 hpi from BJ cells infected with AAV2 alone (gcp/ cell = 20’000) or co-infected with AAV2 (gcp/ cell = 20’000) and HSV-1 (pfu/ cell = 1) showing untreated, *Hin*dIII digested or mung bean nuclease (MBN) treated samples.

### Validation of nanopore sequencing for the detection of an RCR mechanism

To further validate our sequencing approach for determining viral genome replication mechanisms and support our finding that the predominant concatemer structure of HSV-1 assisted AAV2 genome replication are head-to-tail concatemers generated by RCR, we determined the replication products from a well characterized rolling circle amplification (RCA) (29). Specifically, we analyzed the structures generated upon the isothermal bacteriophage Phi29 polymerase mediated RCA of the plasmid pUC19. As shown in **Table 2** and **Supplementary Figure S3**, we found the same structures of parallel full plasmid repeats as described above as category 3. These reads make up 30.5% of all analyzed pUC19 reads. We also found a substantial number (64%) of reads showing a structure with a combination of head-to-tail and alternating concatemers as described for category 5. Phi29 polymerase-mediated generation of ds DNA was previously characterized and suggested to occur upon template switch (30). Template switch of the RCA product would lead to concatemers containing head-to-tail repeats, which change polarity to continue further with head-to-tail repeats as seen in our reads assigned to category 5. Therefore, those previous results are in line with our observations. Furthermore, we found structures resembling the aforementioned duplex structures but to a lesser extent (2%). These results clearly show that Phi29-supported RCA results in the formation of structures similar to those isolated from AAV2 and HSV-1 co-infected cells. Additionally, 3.5% of reads could not be allocated to any of the other specific categories.

**Table 2.** Read analysis data of Phi29 polymerase mediated RCR of pUC19 DNA.

| Category: | 1 | 2 | 3 | 4 | 5 | 7 |  |
| --- | --- | --- | --- | --- | --- | --- | --- |
| Sample | Monomer | Duplex | Head-to-Tail Repeats | Alternating Repeats | Head-to-Tail and Alternating Repeats | Others |  |
|  | ratio | ratio | ratio | ratio | ratio | ratio | total reads |
| pUC19 RCR | 0 | 0.020 | 0.305 | 0 | 0.640 | 0.035 | 200 |

### Analysis of minute virus of mice DNA replication products

To validate our approach further, we analyzed sequences of DNA replication intermediates from the autonomous parvovirus minute virus of mice (MVM). MVM genome replication has been characterized in detail and was described to occur as a modified RHR mechanism similar to the one suggested for AdV-supported AAV genome replication (9, 10, 13, 31). In contrast to AAV, the MVM genome is flanked by two non-identical palindromic terminal (heterotelomeric) sequences and only one of those undergoes site-specific nicking during replication (10, 32–34). As a result, packaged MVM genomes are mainly, although not exclusively, present as ss DNA of negative orientation (1). Furthermore, MVM duplex structures have a preference for a covalently linked (left) 5’-OH-end (31). Extrachromosomal DNA isolated at 16 or 20 hpi from MVM-infected A9 cells, when viral DNA synthesis is maximal and before packaging (35), were sequenced and analyzed as described above. Each read was assigned to one of the previously defined categories (Figure 1b). Two representative dot plots per category are shown in **Figure 4**. Monomeric genome structures were less abundant in MVM-samples (14.5-25.5%) when compared to genomes isolated from AAV2 infected cells (17.5-62%) (**Tables 1 and 3**). However, as mentioned above, quantification of monomeric genome structures should be considered with caution as only ds DNA was described to be sequenced with our approach. Since MVM-genomes are predominantly packaged as negative sense ssDNA, only a small fraction of the genomes will anneal to a positive ss DNA and subsequently be sequenced. The most abundant structure of isolated MVM DNA was the duplex form (67-79.5%) (**Table 3**). MVM was described to replicate its genome via an obligatory dimer replicative form molecule (13) and therefore, our observation of a predominant duplex structure is in line with the textbook model of MVM replication. Those duplex structures were mainly present with a covalent link at the 5’-OH-end (**“V” structure; Supplementary Table S2**). No such preference of duplex structure was found in AAV2-infected cells (**Supplementary Table S2)**. These observations are also in line with the previously described model of MVM genome replication (13, 31). Importantly, no reads containing head-to-tail concatemers (category 3) were identified as MVM-replication intermediates. In contrast, concatemers of alternating repeats were indeed observed, although with a low abundance (0-2.5%). None of the reads showed a combination of head-to-tail and alternating repeats or multiple terminal end-repeats (categories 5 and 6). No clear difference in genome structure between samples harvested at 16 or 20 hpi was observed. However, MVM genome concatemers of alternating repeats were more abundant at the lower multiplicity of infection (1’000 gcp/ cell versus 10’000 gcp/ cell). Overall, we identified replication intermediates as expected from the previously described replication model of the MVM genome (9, 10, 13, 31) and did not find any indication for the generation of RCR-like replication products. This result further validates the use of nanopore sequencing to study viral genome replication mechanisms.

**Table 3.** Read analysis data from extrachromosomal MVM sequences isolated from MVM infected A9 cells.

| Category: |  | 1 | 2 | 3 | 4 | 5 | 6 | 7 |  |
| --- | --- | --- | --- | --- | --- | --- | --- | --- | --- |
| Sample |  | Monomer | Duplex | Head-to-Tail Repeats | Alternating Repeats | Head-to-Tail and Alternating Repeats | Hairpin Repeats | Others |  |
| gcp/cell | harvest | ratio | ratio | ratio | ratio | ratio | ratio | ratio | total reads |
| 1k | 16hpi | 0.225 | 0.730 | 0.000 | 0.025 | 0.000 | 0.000 | 0.020 | 200 |
| 1k | 20hpi | 0.145 | 0.795 | 0.000 | 0.025 | 0.000 | 0.000 | 0.035 | 200 |
| 10k | 16hpi | 0.184 | 0.767 | 0.000 | 0.010 | 0.000 | 0.000 | 0.039 | 103 |
| 10k | 20hpi | 0.255 | 0.670 | 0.000 | 0.000 | 0.000 | 0.000 | 0.075 | 200 |

**Figure 4.**
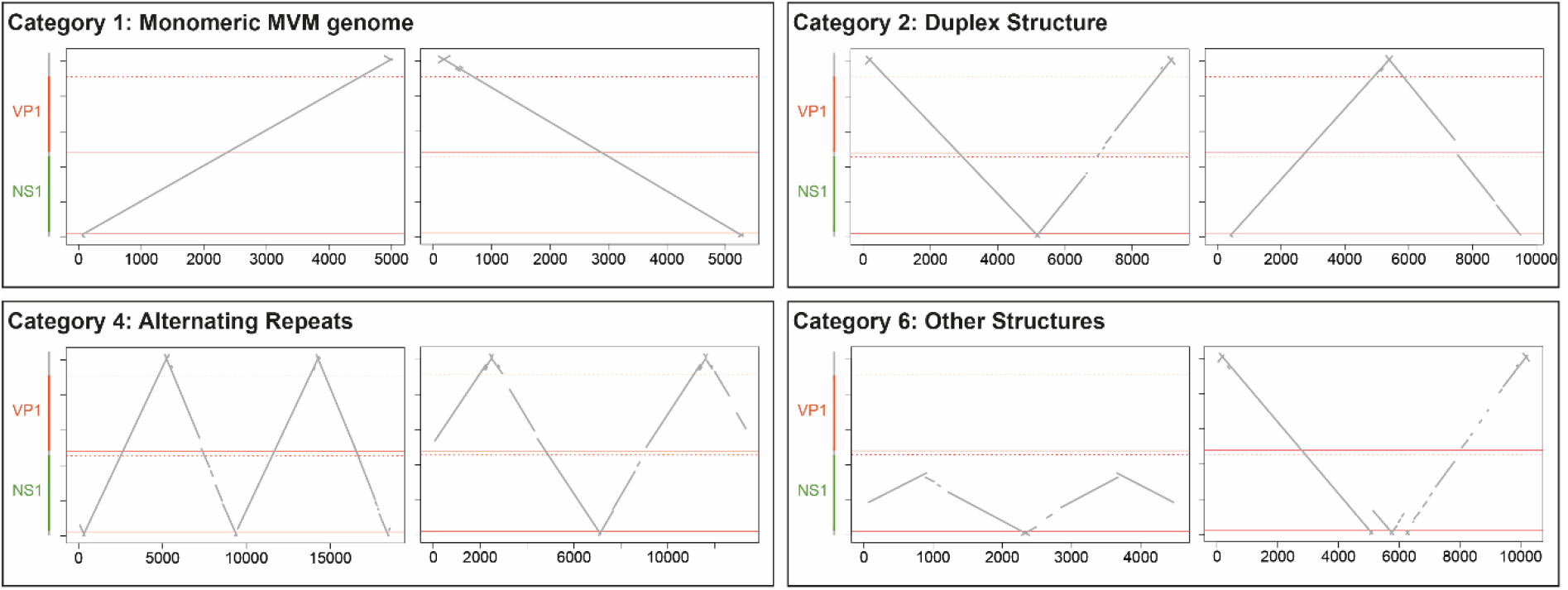
MVM replication intermediates. Dot plots of two representative individual reads per category from MVM infected A9 cells are shown. Extrachromosomal DNA was isolated at 16 hpi from A9 cells infected with MVM (MOI = 1’000).

## DISCUSSION

AAV2 genome replication in the presence of AdV was previously described to occur in a rolling hairpin amplification mode (1, 7, 9, 13). We provide evidence from nanopore sequencing and Southern analysis for an alternative mechanism of AAV2 genome replication when HSV-1 is the helper virus. Our results clearly show the abundant presence of head-to-tail concatemers (category 3), whereas head-to-head concatemers (category 4) were present in very low ratios. The head-to-tail concatemers were neither abundant in AAV2 viral stocks nor in cells infected with AAV2 alone and must therefore be induced by the HSV-1 co-infection. In addition, we identified reads that start as head-to-tail concatemers, then change orientation and continue again as head-to-tail concatemers (category 5). Both categories, 3 and 5, were found in high abundance in Phi29-polymerase mediated RCA samples, supporting our hypothesis that the observed head-to-tail concatemers of the AAV2 genomes arise from an RCR mechanism. However, we cannot exclude an additional RHR in the presence of HSV-1 as we do observe, albeit at very low levels, concatemers of AAV2 genomes with alternating orientation. Also, the low abundance of such concatemers may be due to efficient *trs*-nicking, which prevents concatemer formation.

There are two possibilities of how the head-to-tail concatemers are formed: intermolecular recombination or RCR. We regard RCR as more likely. If head-to-tail concatemers were generated by intermolecular recombination, the probability of concatemers containing multiple genome copies with the same direction would be very low. We observed head-to-tail concatemers of up to 13 repeats in the same direction. The likelihood that such large head-to-tail concatemers were formed upon end recombination would be 0.5^13-1^ = 0.02% (probability of 0.5 for either head-to-head or head-to-tail recombination at each of 12 junctions). Furthermore, if concatemers were generated upon genome end recombination rather than RCR, we would expect more variety in the structure of concatemers and a higher abundance of alternating repeats.

Previous studies identified the helper factors for AAV replication. The essential AdV helper factors include the gene products of *E1a*, *E1b55K*, *E2a*, *E4orf6*, and the VA RNA (36, 37). It was shown that neither the AdV polymerase nor the terminal protein mediate amplification of the AAV2 genome (36, 38, 39). Thus, the AdV helper factors provide indirect support for AAV2 genome replication. The minimal set of HSV-1 helper factors required to support replication in a transient AAV infection model are the helicase-primase complex (HP; UL5/ UL8/ UL52) and the single-strand DNA binding protein ICP8 (40). Although sufficient to promote replication, the HP complex and ICP8 alone were only capable of restoring replication to about 1% compared to wildtype HSV-1 supported replication (41). The addition of plasmids encoding the HSV-1 proteins ICP0, ICP4, ICP22 and the HSV-1 polymerase (UL30/ UL42) together with ICP8 and the HP complex encoding plasmids restored the levels of AAV replication to levels comparable to a wildtype HSV-1 infection (42). The HSV-1 proteins ICP0, ICP4 and ICP22 were found to transactivate AAV *rep* expression, thereby supporting replication (42). While cellular polymerases were found to mediate replication of the AAV genome in the presence of AdV (43, 44), the HSV-1 polymerase was shown to directly support AAV genome replication (44, 45). The essential factors for HSV-1 genome replication comprise ICP8, UL9, the HP complex and the HSV-1 polymerase (46). It was also shown that these factors mediate a rolling circle amplification of non-viral plasmid DNA in the absence of the origin binding protein UL9 (22, 23). Thus, the minimal factors required for AAV genome replication are the same factors that are required to mediate RCR. Since the AAV genome was found to readily circularize even in the absence of a helper virus (25), it is conceivable that the HSV-1 mediated AAV genome replication occurs in an RCR mechanism.

The RCR-mechanism requires a circular template. AAV was found to circularize its genome within the host cell (24, 25) and thus can be used as a substrate during RCR. Next during RCR, the origin of DNA replication is bound by an origin-binding protein and a site-specific nick is introduced into one strand. The large AAV Rep proteins (Rep68/78) were demonstrated to site-specifically bind and nick the AAV genome at the *trs* within the ITRs (19). After nicking occurred, the ds DNA is unwound by a helicase and replicated by a DNA polymerase. Rep68/78 were shown to possess helicase activity (14). Upon completion of an entire amplification cycle, an endonuclease nicks the genome at the origin, releasing the copied DNA. During bacterial or bacteriophage RCR, it is common that the endonuclease, which mediated nicking at the origin, stays covalently attached to the 5’-end possibly mediating the second nicking after replication of the genome (e.g. bacteriophage ϕX147 cistron A protein (47)). Rep68/78 were also demonstrated to remain covalently attached to the 5’-end of the AAV genome (38). Thus, the large AAV Rep proteins possess activities, which are common for RCR-mediating proteins further supporting our hypothesis of an RCR-mechanism during AAV2 replication. The role of Rep68/78 during HSV-1 mediated RCR will be assessed in future studies.

In addition to revealing the structure of not previously described AAV2 DNA replication intermediates, we present a high-throughput method to investigate viral replication mechanisms. The unique feature of a, theoretically, endless sequencing length by the Oxford Nanopore Technology can be used to study replication intermediates on the level of single molecules. In previous approaches, such as Southern analysis, the samples were assessed in bulk and therefore, heterogeneous low-abundant DNA replication intermediates and transient structures were not detected. Our approach revealed the formation of head-to-tail concatemers, unexpected concatemers such as long repeats of the ITR-sequence or a combination of head-to-tail and alternating genome concatemers. Further studies are required to address the question of whether, and how, head-to-tail concatemers can give rise to packageable AAV2 genomes.

## MATERIALS AND METHODS

### Cells and viruses

BJ (foreskin fibroblasts, human), Vero (kidney, African green monkey), Hela (cervical cancer, human), 293T (embryonic kidney, human) and A9 (fibroblasts, mouse) cells were obtained from ATCC (Manassas, Virginia, USA). All cell lines were maintained in Dulbecco’s Modified Eagle’s Medium (DMEM) with high glucose, supplemented with 10% fetal bovine serum (FBS), 2 mM glutamine, 100 units/ml penicillin, 100 μg/ml streptomycin, in a humidified incubator at 37°C and 5% CO_2_. Wildtype HSV-1 (strain F) was grown as previously described (48, 49). A confluent layer of Vero cells was infected at a low multiplicity of infection (MOI) and harvested when cells showed complete cytopathic effect (CPE). Cells were scraped into media, centrifuged, and the cell pellet was frozen and thawed three times. The supernatant and cell pellet were combined and centrifuged again. Aliquots were made from the supernatant and stored at −80°C. Plaque-forming units (pfu) were determined by plaque assay.

Purified AAV2 was produced by the Viral Vector Facility (University of Zurich, Switzerland) as previously described (50, 51). Wildtype AAV2 stocks were produced using the pAV2 plasmid, whereas the AAV2 variant AAV201 was produced using the pSub201 plasmid (52, 53). Stocks of MVM (rodent protoparvovirus 1; clone VR-1346; ATCC) were produced by infection of semi-confluent A9 cell culture as previously described (54). Cell culture supernatant was collected four days post-infection (dpi), cleared by low-speed centrifugation (3000 x g) and stored in aliquots at - 70°C. Viral load was determined by quantitative polymerase chain reaction (qPCR) and median tissue culture infectious dose (TCID50) assay.

### Preparation of AAV2 stock sample for sequencing

The AAV2 stock sample (1.5*10^11^ genome-containing particles) was pre-treated with 10U DNase I (04716728001, Roche, Switzerland) in 1x DNase buffer for 15 min at 37°C to remove any DNA outside the particles. DNase was then inactivated, and capsids were denatured at 95°C for 5 min. The sample was used directly for nanopore sequencing.

### Infection protocol

Cells were seeded 24 h before infection. BJ cells were seeded at 8*10^5^ cells/ tissue culture plate (10 cm in diameter; 172958, Nunc, Thermo Fisher Scientific, Waltham, MA, USA) with 8 tissue culture plates per condition. Virus (AAV2 in the presence or absence of HSV-1) was diluted in an appropriate volume of DMEM (0% FBS). Low infection volumes were chosen to enhance infection but ensuring the complete submersion of the cell layer. Cells were placed in a humidified incubator at 37°C and 5% CO_2_. One to two hours post infection (hpi), the supernatant was removed and sufficient DMEM containing 2% FBS was added. Cells were incubated in a humidified incubator at 37°C and 5% CO_2_ for the indicated time. For MVM infection, A9 cells were inoculated at a confluency of 20-30% to ensure exponential cell growth during replication (55). One T-150 flask per condition was infected with either 10 ml undiluted or 1/10 diluted MVM stock to obtain 10’000 and 1’000 genome equivalents per cell, respectively. The inoculum was removed 1 hpi and replaced with 15 ml DMEM, 10% FBS. Infected cells were incubated in a humidified incubator at 37°C and 5% CO_2_ for the indicated time.

### Hirt extraction

Extraction of extrachromosomal DNA was performed according to the Hirt protocol (56). Cells were washed with PBS and detached using 0.05% Trypsin-EDTA. The cell pellet was resuspended in 50 μl TBS (50 mM Tris-HCl, 150 mM NaCl, pH 7.5). Then 500 μl Hirt buffer (0.6% SDS, 10 mM Tris-HCl, 10 mM EDTA, pH 7.5) was added, and the suspension was incubated at room temperature for 1 h. After adding 120 μl of a 5 M NaCl solution, the sample was incubated at 4°C for at least 12 h. For phenol/ chloroform extraction of DNA, the sample was centrifuged at 15’500 g for 10 min at 4°C. The supernatant was transferred into a fresh tube and 1 volume of phenol: chloroform: isoamyl alcohol (25:24:1, v/v) was added. The sample was centrifuged at 15’500 g for 5 min at 4°C. The supernatant was transferred into a fresh tube and 1 volume of chloroform was added. The sample was centrifuged for 1min at 15’500 g at 4°C. The supernatant was transferred into a fresh tube and 2.5 volumes of EtOH (pure) and 0.1 volume of 3 M NaAc pH 5.5 was added, and the suspension was incubated at −80°C for at least 20 min to precipitate the DNA. The sample was centrifuged at 18’000 g for 10 min at 4°C and the supernatant was discarded. The pellet was washed with 70% EtOH. After centrifugation at 18’000 g for 10 min at 4°C, the supernatant was removed and the pellet was air-dried before being resuspended in 10 mM Tris-HCl, pH 8.5.

### Rolling circle amplification of pUC19

Plasmid DNA (pUC19, 2 ng) was amplified using the circular DNA amplification kit TempliPhi (25-6400-10, Cytiva, Marlborough, MA, USA) according to the manufacturer’s protocol. Then, phenol/ chloroform DNA purification was performed as described in the Hirt extraction protocol above.

### Nanopore sequencing

Nanopore sequencing was performed by the research group of Dr. A. Ramette (University of Bern, Switzerland). Hirt-extracted DNA samples were prepared for sequencing using the ligation sequencing kit (LSK109, Oxford Nanopore Technologies, Oxford, UK) and the native barcoding expansion kit (EXP-NBD104, Oxford Nanopore Technologies, Oxford, UK) according to the manufacturer’s protocol. Subsequently, the samples were sequenced using a GridION X5 sequencing device and flow cell (FLO-MIN-106, Oxford Nanopore Technologies, Oxford, UK).

### Data analysis

Sequences in fastq-format were transformed to fasta-format by seqkit (57). The reads were subjected to blastn (58) analysis against the AAV2 or the MVM genomic sequence (Genbank # NC_001401 or NC_001510) or against the pUC19 plasmid sequence (Genbank #M77789). Lines with no hits and lines with comments were removed (grep -v ‘#’) and tables were read into R 4.0 (The R Foundation for Statistical Computing). Hits with e-values > 0.1 were removed. The sums of all hits were calculated for every read. The reads with a sum of hits < 3000 were removed. Dot plots of the remaining reads were drawn. The dot plots were manually categorized. The code for the bioinformatic analysis can be found in the **Supplementary Material**.

### Southern blot

For Southern analysis, extrachromosomal DNA was extracted using the Hirt protocol. *Hin*dIII treatment was performed at 37°C for 1 h in 1x reaction buffer B with 10U of *Hin*dIII enzyme per reaction (*Hin*dIII, 10656321001, Roche, Switzerland). Mung bean nuclease treatment was performed at 30°C for 30 min in 1x mung bean nuclease reaction buffer with 1U per reaction (Mung Bean Nuclease, M0250S, New England Biolabs, Ipswich, MA, USA). DNA (40 ng per sample) was separated on 0.8% agarose gels and transferred onto nylon membranes (Hybond-N+, RPN119B, Amersham, Little Chalfont, UK). As reference, DIG-labeled marker was used (DNA molecular weight marker III, 11218602910, Roche). Hybridization with a digoxigenin (DIG)-labeled probe specific for AAV2 *rep,* subsequent detection by an anti-DIG antibody conjugated with alkaline phosphatase and activation with the chemiluminescence substrate CDP Star (Roche) were performed according to the manufacturer’s protocol. The DIG-labeled probe was synthesized using the PCR DIG probe synthesis kit (11636090910, Roche, Switzerland) and the following primers: 5’-gaa cgc gat atc gca gcc gcc atg ccg gg-3’ and 5’-gga tcc gaa ttc act gct tct ccg agg taa tc-3’. Chemiluminescence was imaged using the LI-COR imaging system Odyssey Fc (LI-COR Biosciences, Lincoln, NE, USA).

## Data Availability

Raw sequencing data used in this study have been deposited at the National Center for Biotechnology Information Sequence Read Archive (NCBI SRA) website (https://www.ncbi.nlm.nih.gov/sra) under the accession numbers PRJNA699763 and PRJNA699846.

## ACKNOWLEDGEMENTS

We thank Mathias Ackermann for the scientific discussions. We thank Alban Ramette and Stefan Neuenschwander for their support with nanopore sequencing. We thank Dinis Meier for his support in establishing the design of the figures. We thank Tobias Myland for proofreading the manuscript. This work was funded by the Swiss National Science Foundation No. 310030_184766 to C.F.

