## Supplementary for "Herpes Simplex Virus Co-Infection Facilitates Rolling Circle Replication of the Adeno-Associated Virus Genome"

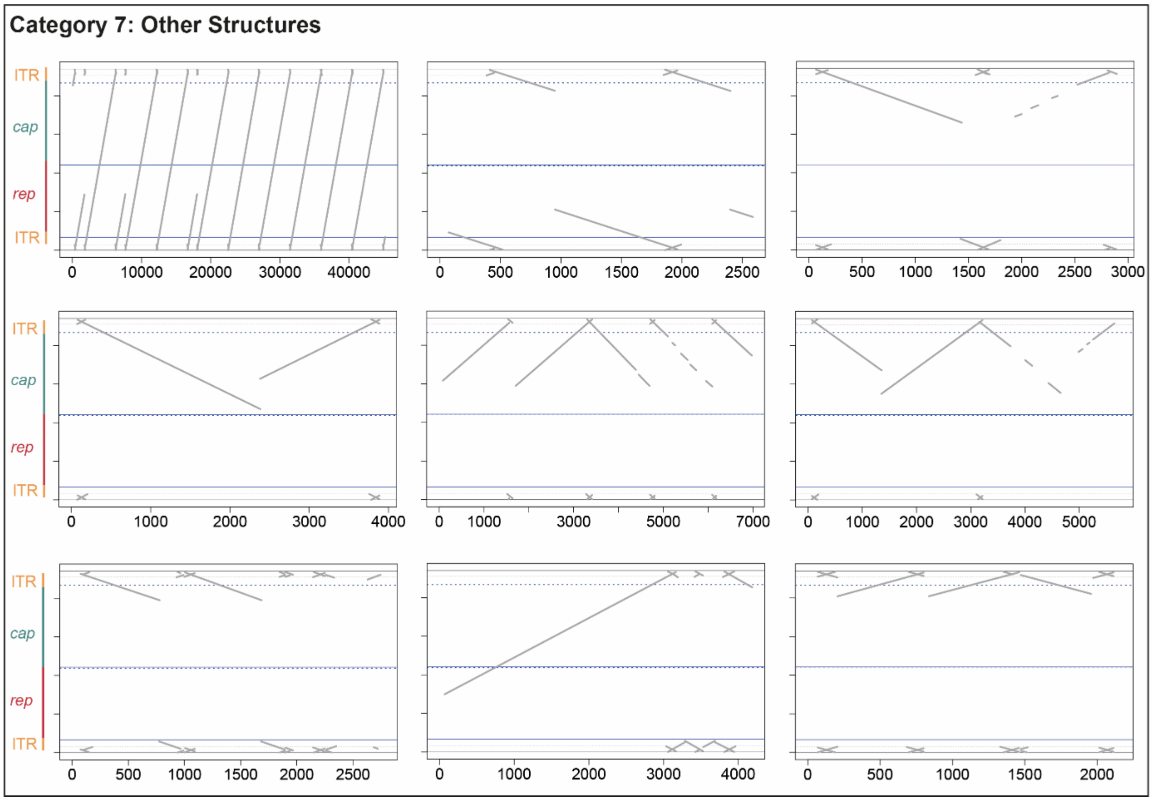


**Figure S1.** AAV2 DNA replication intermediates. Dot plots of individual reads assigned to category 7 (other structures) from AAV2/ HSV-1 co-infected BJ cells are shown. Extrachromosomal DNA was isolated at 12 hpi from BJ cells infected with AAV2 (gcp/ cell = 20’000) and HSV-1 (pfu/ cell = 1).

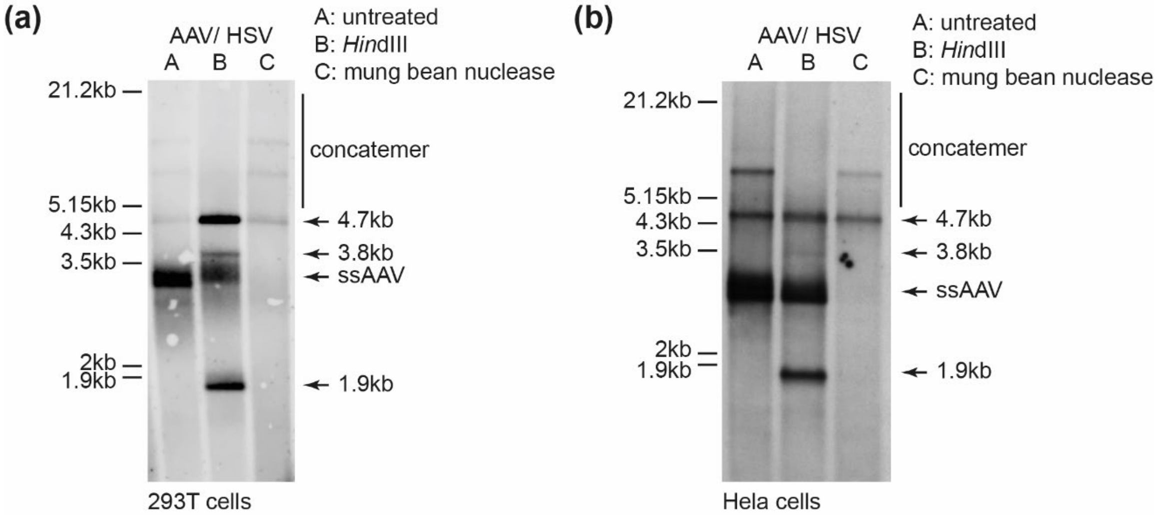


**Figure S2.** Southern analysis of AAV2 DNA. Southern blot of Hirt DNA extracted at 15 hpi from 293T cells (a) or at 20 hpi from Hela cells (b) co-infected with AAV2 (gcp/ cell = 20'000) and HSV-1 (pfu/ cell = 1) showing untreated (A), *Hin*dIII digested (B) or mung bean nuclease treated (C) samples.

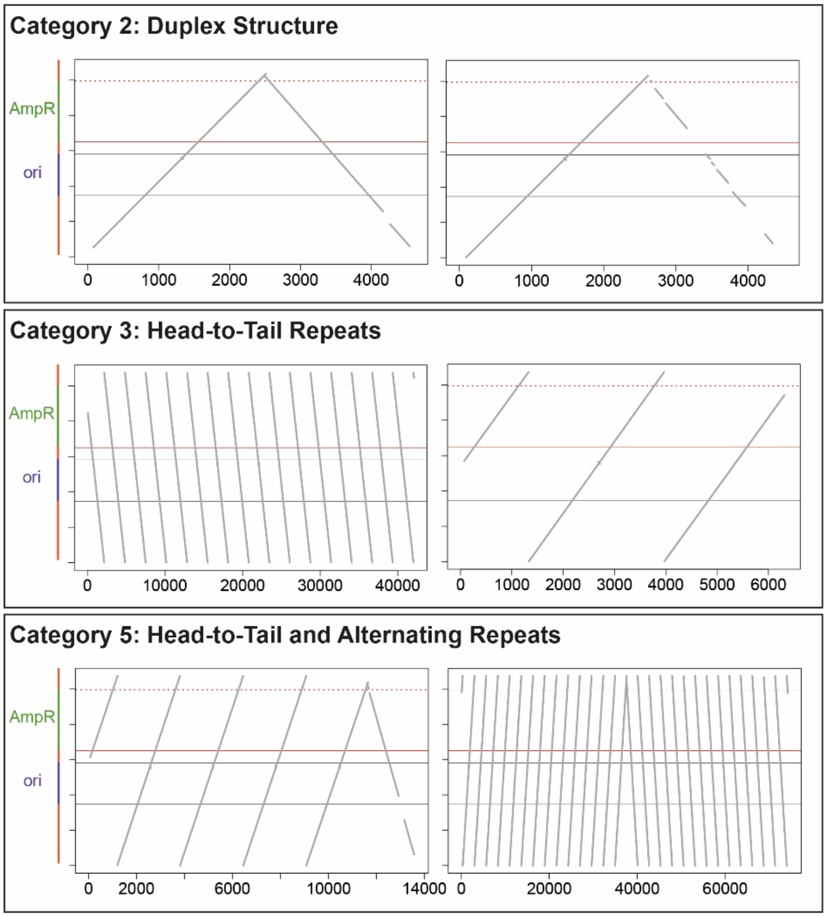


**Figure S3.** Amplification products of Phi29 polymerase mediated RCA on pUC19. Dot plots of two individual reads per category of DNA isolated from Phi29 polymerase amplified pUC19.

**Table S1.** Read analysis data from genomes isolated from AAV201 single- or HSV-1 co-infected BJ cells at 12 hpi.

| Category: | | 1 | 2 | 3 | 4 | 5 | 6 | 7 | |
|---|---|---|---|---|---|---|---|---|---|
| Sample | | Monomer | Duplex | Head-to-Tail Repeats | Alternating Repeats | Head-to-Tail and Alternating Repeats | ITR Repeats | Others | |
| | AAV gcp/ cell | ratio | ratio | ratio | ratio | ratio | ratio | ratio | total reads |
| **AAV201** | **20k** | 0.632 | 0.098 | 0.000 | 0.000 | 0.000 | 0.127 | 0.142 | 204 |
| **AAV201 / HSV-1** | **20k** | 0.162 | 0.221 | 0.235 | 0.025 | 0.054 | 0.108 | 0.196 | 204 |

**Table S2.** Further read analysis data from extrachromosomal MVM sequences isolated from MVM infected A9 cells or AAV2 sequences from AAV2 single- or HSV-1 co-infected BJ cells of category 2.

| Category: | | | 2 | 2a | 2b | |
|---|---|---|---|---|---|---|
| Sample | | | 2 Duplex Structure (total) | 2a Duplex Structure "A" | 2b Duplex Structure "V" | |
| **Infection** | **MOI** | **harvest** | **ratio** | **ratio** | **ratio** | **total reads** |
| **MVM** | **1k** | **16hpi** | 0.730 | 0.050 | 0.680 | 200 |
| **MVM** | **1k** | **20hpi** | 0.795 | 0.065 | 0.730 | 200 |
| **MVM** | **10k** | **16hpi** | 0.767 | 0.087 | 0.680 | 103 |
| **MVM** | **10k** | **20hpi** | 0.670 | 0.100 | 0.570 | 200 |
| **AAV2** | **20k** | **12hpi** | 0.080 | 0.040 | 0.040 | 200 |
| **AAV2/ HSV-1** | **20k/ 1** | **12hpi** | 0.170 | 0.105 | 0.065 | 200 |
| **AAV2** | **500** | **12hpi** | 0.333 | 0.083 | 0.250 | 12 |
| **AAV2/ HSV-1** | **500/ 1** | **12hpi** | 0.385 | 0.095 | 0.290 | 200 |

**Bioinformatic Code**

####################################################################################
##### on UNIX platform on UNIX platform on UNIX platform on UNIX platform on UNIX platform on UNIX ###
####################################################################################

Change nanopore reads from fastq- to fasta-format using seqkit:
[W Shen, S Le, Y Li, F Hu. "SeqKit: a cross-platform and ultrafast toolkit for FASTA/Q file manipulation." PLOS ONE. doi:10.1371/journal.pone.0163962.] download and install: https://bioinf.shenwei.me/seqkit/download/

### execute command:
```
seqkit fq2fa <path_to_fastq_files.fastq> -o <path_to_fasta_files.fasta>
```

Alignment of reads in fasta-format with blastn against ref_seq:
[Christiam Camacho 1 , George Coulouris, Vahram Avagyan, Ning Ma, Jason Papadopoulos, Kevin Bealer, Thomas L Madden "BLAST+: architecture and applications" BMC Bioinformatics doi: 10.1186/1471-2105-10-421.] download and install: https://ftp.ncbi.nlm.nih.gov/blast/executables/blast+/LATEST/

### execute command:
```
blastn -word_size 11 -reward 2 -penalty -3 -query <path_to_fastq_files.fasta> -
db <path_to_blastn_db> -outfmt "7 qacc sacc evalue qstart qend sstart send
qlen" -out <path_to_outfile>
```

### remove lines with non-hits and comments from blastn-output and save to text-file:
```
grep -v '#' <path_to_outfile> > <path_to_outfile.txt>
```

####################################################################################
### in R in R in R in R in R in R in R in R in R in R in R in R in R in R in R in R in R in R in R in R in R in R in R ###
####################################################################################

Read values of blastn hit table into R and store as dataframe ‘sample_name’:
```
sample_name <- read.table("<path_to_outfile.txt>", sep="\t",
col.names=c("read_ID", "sub_name", "e_val", "que_beg", "que_end", "sub_beg",
"sub_end", "que_length"))
```

Reduce dataframe for hits with e-value smaller than 0.1:
```
sample_name <- sample_name[sample_name$e_val < 0.1, ]
```

Order dataframe according to the read_ID:
```
sample_name <- sample_name[order(sample_name$read_ID), ]
```

Generate column in dataframe with rep_hits:

```
#######################################################################################
# define function to generate vector w/ identical read nummber for all hits of same read:
# USAGE: df$rep_hits <- rep.hits(df)

rep.hits <- function(df) {
x <- c(); n <- tabulate(as.factor(df$read_ID));
for (i in 1:length(n)) {x <- append(x, rep(i, n[i]))}

return(x)
}
#======================================================================================
```

Apply rep.hits-function on dataframe:

```
sample_name$rep_hits <- rep.hits(sample_name)
```

Generate column in dataframe with tot_hits:

```
########################################################################################
# define function to generate vector w/ total hit length for each read:
# USAGE: df$tot_hits <- tot.hits(df)

tot.hits <- function (df) {

sum_read <- c(); sum_vect <- c();
for (i in 1:max(df$rep_hits)) {
                  sum_read <- sum(df[df$rep_hits == i,]$que_end - df[df$rep_hits ==
i,]$que_beg);
                  sum_vect <- append(sum_vect, rep(sum_read, tabulate(df$rep_hits)[i]));
                             }
return(sum_vect);
                          }
#=======================================================================================
```

Apply tot.hits-function on dataframe:

```
sample_name$tot_hits <- tot.hits(sample_name)
```

Reduce dataframe for tot_hits larger than 3000nt:

```
sample_name <- sample_name[sample_name$tot_hits > 3000, ]
```

Generate NEW rep_hits column in dataframe:

```
sample_name$rep_hits <- rep.hits(sample_name)
```

Define functions to draw dotplots:

```
#########################################################################################
# define function to draw one dotplot of "wtAAV" blast hits from one nanopore read:
# USAGE: dot.wt(df)

dot.wt <- function (df) {

plot(NULL, xlim=c(0, df$que_length[1]), ylim=c(0, 4700), xlab="Query", ylab="Subject",
main=df$read_ID[1]);
# draw annotation on dt plot:
abline(h=329, col="blue"); abline(h=2194, lty=2, col="blue"); abline(h=2211,col="blue");
abline(h=4337, lty=2, col="blue");
text(1,1261,"rep", cex=0.5, col="blue"); text(1,3274,"cap", cex=0.5,col="blue")
abline(h=0, lwd=0.5); abline(h=150, lty=2, lwd=0.5); abline(h=4550, lty=2, lwd=0.5);
abline(h=4700, lwd=0.5);
text(1,50,"ITR", cex=0.5); text(1,4600,"ITR", cex=0.5)

# draw individual hits:
for (i in 1:length(df$read_ID)) {
segments(df$que_beg[i], df$sub_beg[i], df$que_end[i], df$sub_end[i], col="darkgrey",
lwd=3);
                                }
                           }
#========================================================================================

#########################################################################################
# define function to draw dotplots blast hits on all nanopore reads from one sample:
# USAGE: dot.all(df, dot.wt)

dot.all <- function (df, func) {

for (j in 1:max(df$rep_hits)) {
                  temp_df <- df[df$rep_hits == j,];
                  func(temp_df);
                                }
                               }
#========================================================================================
```

Generation of dotplots from all nanopore reads redirected into a pdf-file:

```
pdf("<path_to_output.pdf>", paper="a4", width=7.3, height=10.7)
par(mfrow=c(4,2))
dot.all(sample_name, dot.wt)
dev.off()
```
